# Time-Invariant Working Memory Representations in the Presence of Code-Morphing in the Lateral Prefrontal Cortex

**DOI:** 10.1101/563668

**Authors:** Aishwarya Parthasarathy, Cheng Tang, Roger Herikstad, Loong Fah Cheong, Shih-Cheng Yen, Camilo Libedinsky

**Author notes:** Shih-Cheng Yen and Camilo Libedinsky contributed equally to this work.

## Abstract

Endogenous processes allow the maintenance of working memories. These processes presumably involve prefrontal networks with strong recurrent connections. Distractors evoke a morphing of the population code, even when memories are stable. But it is unclear whether these dynamic population responses contain stable memory information. Here we show that dynamic prefrontal activity contains stable memory information, and the stability depends on parallel movement of trajectories associated with different memories in state space. We used an optimization algorithm to find a subspace with stable memory information. In correct trials the stability extended to periods that were not used to find the subspace, but in error trials the information and the stability were reduced. A bump attractor model was able to replicate these behaviors. The model provided predictions that could be confirmed with the neural data. We conclude that downstream regions could read memory information from a stable subspace.

## Introduction

Working memory (WM) is the ability to hold and manipulate information over a short time. It is a core component of complex cognitive functions, such as reasoning and language (Baddeley 2012). Memory maintenance appears to involve the sustained activity of neurons in the lateral prefrontal cortex (LPFC) (Constantinidis et al., 2018; but see Lundqvist et al., 2018), and populations of LPFC neurons have exhibited time-invariant (stable) codes during the memory-maintenance period (Stokes et al., 2013; Mendoza-Halliday et al., 2017). Distractors presented during WM maintenance disrupt code stability in the LPFC, despite behavioral evidence of WM stability (Parthasarathy et al., 2017; Cavanagh et al., 2018). If the LPFC plays a role in WM maintenance, then stable memory information should be present in the morphing code.

We hypothesized that a LPFC response subspace retains stable memory information in the presence of code-morphing. We used an optimization algorithm that minimized the subspace distance between the Delay 1 and Delay 2 responses using a cost function that included a penalty for information loss. Using this optimization, we found a subspace with a stable memory code. The stability extended to the distractor presentation period, which was not used in the optimization, and the stability was absent in error trials. These results show that the LPFC retains behaviorally relevant stable memory information despite exhibiting code-morphing.

Network models have been shown to replicate several properties of LPFC activity, including code stability (Compte et al., 2000; Wimmer et al., 2014; Murray et al., 2017; Yang et al., 2019). However, it is not known whether these models can exhibit code-morphing while retaining a stable subspace. We found that a bump attractor model with different target and distractor inputs was most effective in replicating the results we observed. These results suggest that non-memory inputs to the LPFC may be a critical component of code-morphing.

## Results

Two adult monkeys were trained to perform a delayed saccade task with an intervening distractor (Fig. 1a). We recorded a total of 256 neurons from the LPFC and 137 neurons from the FEF while the animals performed the task. Cross-temporal decoding (Fig. 1b) and state-space analysis (Fig. 1c) showed that the distractor presentation led to code-morphing in the LPFC (quantified in Fig. 2d,e), as previously described (Parthasarathy et al., 2017, see Supplementary Movie 1 for an illustration of the trajectories).

**Fig. 1.**
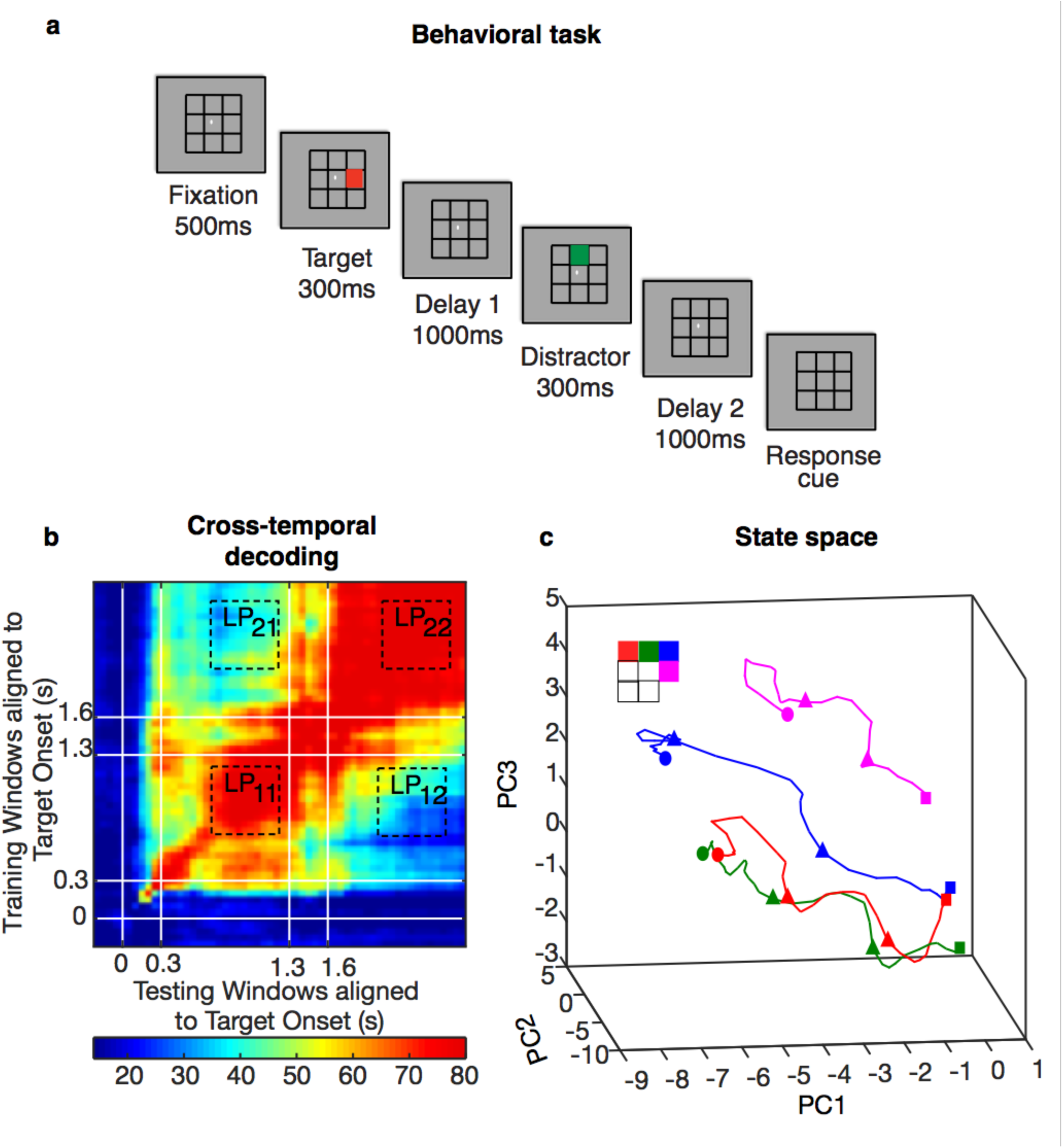
Experimental design and code-morphing. **a**, Behavioral task: Each trial began when the animal fixated on a fixation spot in the centre of the screen. The animal was required to maintain fixation throughout the trial until the fixation spot disappeared. A target (red square) was presented for 300 ms followed by a 1000-ms delay period (Delay 1). A distractor (green square) was then presented for 300 ms in a random location that was different from the target location, and was followed by a second delay of 1000 ms (Delay 2). After Delay 2, the fixation spot disappeared, which was the Go cue for the animal to report, using an eye movement, the location of the target. **b**, Heat map showing the cross-temporal population-decoding performance in the LPFC. White lines indicate target presentation (0–0.3 s) and distractor presentation (1.3–1.6 s). **c**, Responses of the LPFC population when projected onto the first three principal components (PC) of the combined Delay 1 and Delay 2 response space. The responses for different target locations are color-coded using the color scheme shown in the top left. The trajectories illustrate the evolution of the responses from 500 ms before the end of Delay 1 (square), distractor onset (first triangle), distractor offset (second triangle), and the end of Delay 2 (circle). The trajectories of only 4 of the 7 target locations are shown here for clarity.

**Fig. 2.**
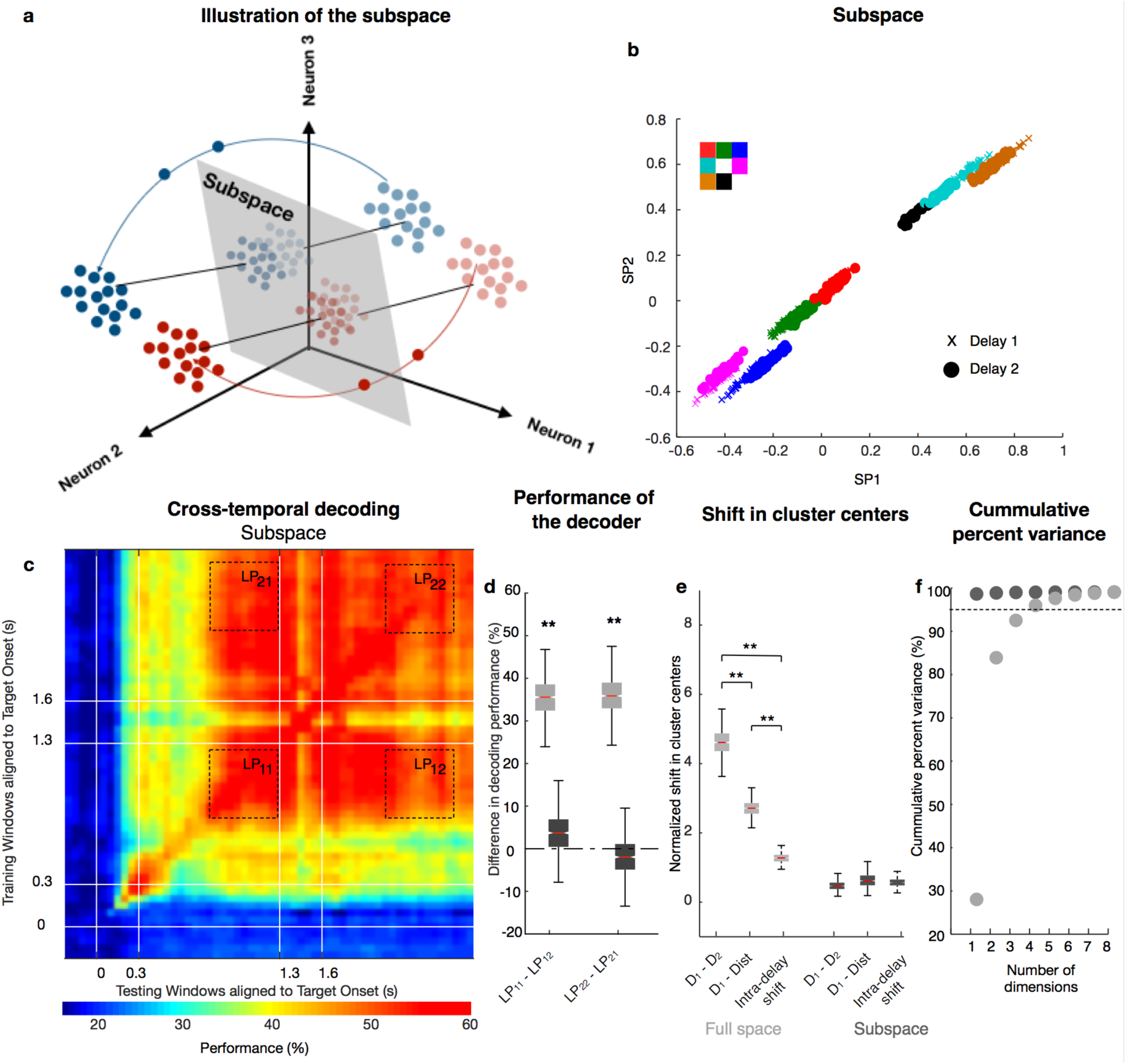
Identification of a subspace. **a**, Illustration of the subspace where Delay 1 and Delay 2 activity exists as one persistent code. **b**, Delay 1 (plotted using crosses) and Delay 2 (plotted using squares) responses after projection into the top 2 PCs of the subspace. Points for different target locations are color-coded according to the color scheme shown in the top left. **c**, Heat map showing the cross-temporal population-decoding performance after the population responses were projected onto the subspace. White lines indicate target presentation (0–0.3 s) and distractor presentation (1.3–1.6 s). **d**, The difference in decoding performance when a decoder that was trained on Delay 1 responses was tested on Delay 1 (*LP*_*11*_) or Delay 2 (*LP*_*12*_) responses are shown in the gray box-plot labelled *LP*_*11*_ - *LP*_*12*_, and vice versa (gray box-plot labelled *LP*_*22*_ - *LP*_*21*_). The equivalent performance differences in the subspace are shown in the black box-plots. **e**, The shift in cluster centers from Delay 1 to Delay 2 (labeled D1 - D2) averaged across target locations, from Delay 1 to the distractor presentation period (D1 - Dist), and the intra-delay shifts in both Delays 1 and 2 (intra-delay) are shown in the full space (gray box-plot), and in the subspace (black box-plot). **f**, The cumulative explained variance is plotted as a function of the number of PCs for the full space (plotted in gray) and the subspace (plotted in black). In all panels, asterisks (**) denote significance (i.e., 95th percentile range of the distribution did not overlap with zero, or the 95th percentile range of two distributions did not overlap).

The presence of code-morphing presents an interesting decoding challenge: how can downstream regions read out stable information from a morphing code? We hypothesized that there may be a low-dimensional subspace embedded in the LPFC population response that contained stable memory information (Fig. 2a) that could be used by downstream cells. We used an optimization algorithm that minimized the distance between Delay 1 and Delay 2 responses when projected into a reference subspace, while simultaneously maintaining memory information (see Methods). The result of the optimization is shown in Fig. 2b, which shows overlapping Delay 1 and Delay 2 projections in the subspace. As hypothesized, classifiers trained in this subspace at one time point in either of the two delay periods were able to decode memory information equally well from other time points during both delay periods (Fig. 2c). In order to quantify the stability of the code in the subspace, we calculated the difference in decoding performance between decoders trained and tested in Delay 1 (LP11), and decoders trained in Delay 1 and tested in Delay 2 (LP12). Similarly, we compared the performance difference for decoders trained and tested in Delay 2 (LP22), and decoders trained in Delay 2 and tested in Delay 1 (LP21). In this analysis, values above zero implied code-morphing, while values around zero implied a stable code. For decoders built using the full space, we found significant differences, consistent with code-morphing (Fig. 2d, light gray, P < 0.001 for LP11 - LP12 and LP22 - LP21). On the other hand, for decoders built using the subspace, we found no difference, consistent with a stable code (Fig. 2d, dark gray, P ≈ 0.22 for LP11 - LP12 and P ≈ 0.66 for LP22 - LP21).

We also quantified the stability of the code using state space analysis by calculating the mean shift in cluster centers between Delay 1 and Delay 2 (inter-delay shift), compared to the mean intra-delay shift in Delays 1 and 2 (Fig. 2e). In this analysis, inter-delay shifts that were larger than the intra-delay shifts implied code-morphing, while similar shifts implied a stable code. In the full space (Fig. 2e, light gray), we found significantly larger inter-delay shifts when compared to intra-delay shifts (P < 0.001, g = 7.40), consistent with code-morphing. On the other hand, in the subspace (Fig. 2e, dark gray), no such differences were found (P ≈ 0.97), consistent with a stable code.

In order to determine the dimensionality of the full space and the subspace, we computed the cumulative percent variance explained by different numbers of PCA components (Fig. 2f). In the full space, the top four components were required to explain at least 95% of the variance, while in the subspace, only the first component was required to explain at least 95% of the variance. This supported the view that the subspace was clearly different from the full space, and existed in a lower dimensional space within the full state-space.

The optimization algorithm used in this study to identify the stable subspace used activity from the last 500 ms of Delays 1 and 2. We selected these periods because both exhibited internal stability. To our surprise, the subspace stability extended to periods that were not used in the optimization, namely distractor period (P ≈ 0.78 for decoding performance difference; P ≈ 0.96 for cluster shifts) and the first 500 ms of Delay 2 (P ≈ 0.45 for decoding performance difference; Fig 2c,e). The stability of the subspace, however, did not extend to the target presentation period (P < 0.001 for decoding performance difference), nor the first 500 ms of Delay 1 (P < 0.001 for decoding performance difference; Fig. 2c). This observation may reflect a shortcoming of our method in identifying the true subspace that was used from the moment of stimulus presentation until the response. Alternatively, it may reflect a real specialization of the LPFC in encoding memories approximately 800 ms after target presentation (e.g. Murray et al., 2017, Mendoza-Halliday et al., 2017, Cavanagh et al., 2018), which may be the time when activity in the subspace we identified started encoding memory information (Fig. 2c).

Code-morphing depended on the activity of neurons with mixed selectivity (Rigotti et al., 2013), which in our task were defined as those that exhibited simultaneous selectivity to multiple task parameters, such as memory location and task epoch (Parthasarathy et al., 2017). Of particular relevance for code-morphing appeared to be those neurons with non-linear mixed selectivity (NMS), which exhibited an interaction between the selectivity to target location and task epoch. In other words, NMS neurons had different selectivity before and after the distractor (Parthasarathy et al., 2017). A fraction of the neurons in LPFC were classically selective, meaning that they were only selective to memory location or task epoch, but not both, and did not change their selectivity after the distractor. One possible interpretation of the subspace could be that it was built from the activity of classically selective neurons, which by definition would have a stable code. While this would be a simple explanation for the existence of the subspace, it was unlikely, since classically-selective cells contained comparatively little information about the target location, likely due to their poor selectivity (Supplementary Fig. 1a-c). Furthermore, individual neuron contributions to the subspace were similarly distributed across NMS, classically-selective, and linear mixed selective cells (Supplementary Fig. 1d). A subspace could also be identified in a population exclusively composed of NMS neurons, highlighting that the classically-selective neurons were not essential to the formation and maintenance of stable memory information (Supplementary Fig. 2).

The results discussed so far showed that stable memory information could be extracted from a subspace of LPFC neurons exhibiting code morphing. In order to understand whether this finding reflected a coding property of LPFC neurons, or whether any random set of state-space trajectories contain a subspace with stable memory information, we carried out the same analysis on data where the memory locations in Delay 2 were shuffled with respect to Delay 1 (Fig. 3a,b). Using the shuffled data, we were not able to find a subspace that provided a stable memory readout (Fig. 3c,d). The optimization procedure yielded a subspace with low information content (Fig. 3d), and which still displayed code-morphing (Fig. 3e,f; LP11 - LP12: P < 0.001; LP22 - LP21: P < 0.001; shift in cluster centers: true data versus intra-delay shift: P ≈ 0.96; shuffled data versus intra-delay shift: P < 0.04, g = 3.04). These results imply that the existence of a subspace with stable memory information reflects a non-trivial organizational property of LPFC activity. In order to characterize this organizational property we employed a dynamical systems approach, by analyzing the trajectory dynamics of population activity in state space (Remington et al., 2018). For the subspaces created using true and shuffled data, we calculated the average trajectory directions, which is the magnitude of the vector obtained by averaging the trajectory vectors from Delay 1 to Delay 2 for each target location. In this analysis, low values of trajectory direction would be expected from trajectories that move in non-parallel ways. We found that trajectories in the subspace built using true data moved in significantly more parallel trajectories than in the subspace built using shuffled data (Fig. 3g, P < 0.001, g = 1.74). This suggested that the parallel movement of trajectories could be an important response property of LPFC activity that facilitated the existence of a subspace with stable memory information.

**Fig. 3.**
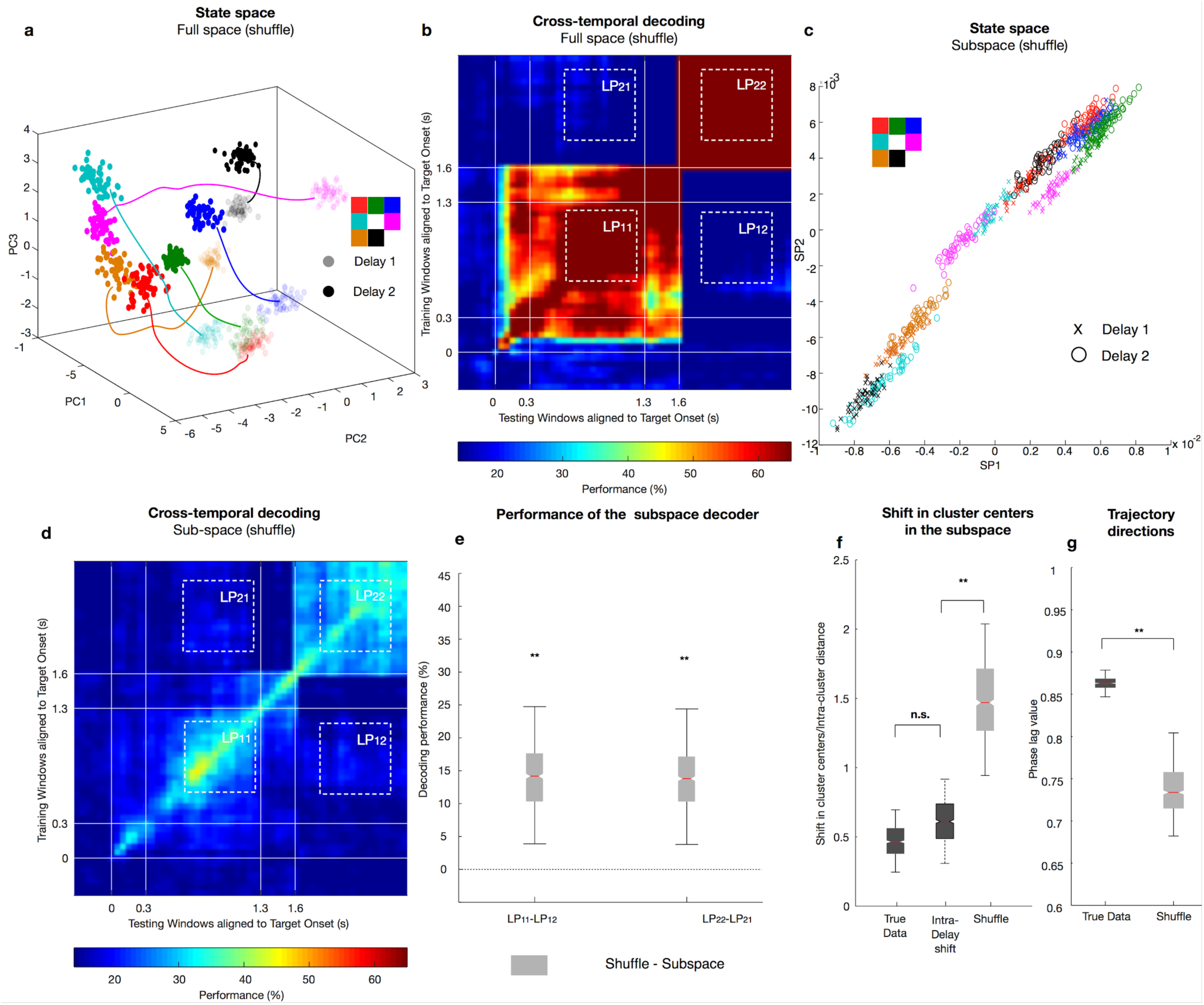
Comparison of the subspace to a shuffled subspace. **a**, State space showing the target locations in Delay 2 (dark colored dots) were shuffled relative to those in Delay 1 (light colored dots) before the optimization was performed in the full space. **b**, Heat map showing the cross-temporal population-decoding performance in the full space with the target location in Delay 2 shuffled relative to those in Delay 1. **c**, Delay 1 (plotted using crosses) and shuffled Delay 2 (plotted using open circles) responses after projection into the top 2 PCs of the subspace. Points for different target locations are color-coded according to the color scheme shown in the top left. **d**, Heat map showing the cross-temporal population-decoding performance in the subspace with the target locations in Delay 2 shuffled relative to those in Delay 1. **e**, The difference in decoding performance when a decoder that was trained on the Delay 1 responses shown in (c) was tested on Delay 1 (LP11) or Delay 2 (LP12) responses are shown in the gray box-plot labelled LP11 - LP12, and vice versa (labelled LP22 - LP21). The equivalent performance differences for the decoding performance in the subspace previously shown in Fig. 2d are shown here in the white boxplots. **f**, The shift in cluster centers from Delay 1 to Delay 2 in the subspace, normalized to the average intra-delay distance, is shown for the true data (black box-plot) and the shuffled data (gray box-plot). The boxplot in the middle illustrates the intra-delay shifts for comparison. **g**, The magnitude of the vector obtained by averaging the trajectory vectors from Delay 1 to Delay 2 for each target location is shown for the true data (black box-plot) and the shuffled data (gray box-plot). Asterisks (**), significant (i.e., 95th percentile range of two distributions did not overlap); n.s., non-significant (i.e., 95th percentile range of two distributions overlapped with each other).

Although we were able to identify a subspace in which the target information could be stably decoded throughout Delays 1 and 2 in spite of code-morphing, it was not clear whether this subspace was related to the behavioral performance of the animals in the task. We investigated this question by comparing the responses of correct and incorrect trials in the subspace. The cross-temporal decoding performance for error trials in the subspace is shown in Figure 4a. Compared to the performance for correct trials shown in Figure 2c, the error trials clearly exhibited much lower performance. We found the performance difference between correct and incorrect trials in the subspace was not significantly different from that in the full space (Fig. 4b) for both Delays 1 (P ≈ 0.74) and 2 (P ≈ 0.25). By analyzing the shift in cluster centers in the subspace, we found that error trials began to deviate from correct trials in Delay 1 (although the deviation did not reach statistical significance, P ≈ 0.06), and become significantly different from correct trials in Delay 2 (Fig. 4c, P ≈ 0.03, g = 2.98). Finally, in error trials, the code stability is disrupted within the subspace, leading to code morphing (Fig. 4d). These results show that behavioral errors were associated with decreases of information within the subspace.

**Fig. 4.**
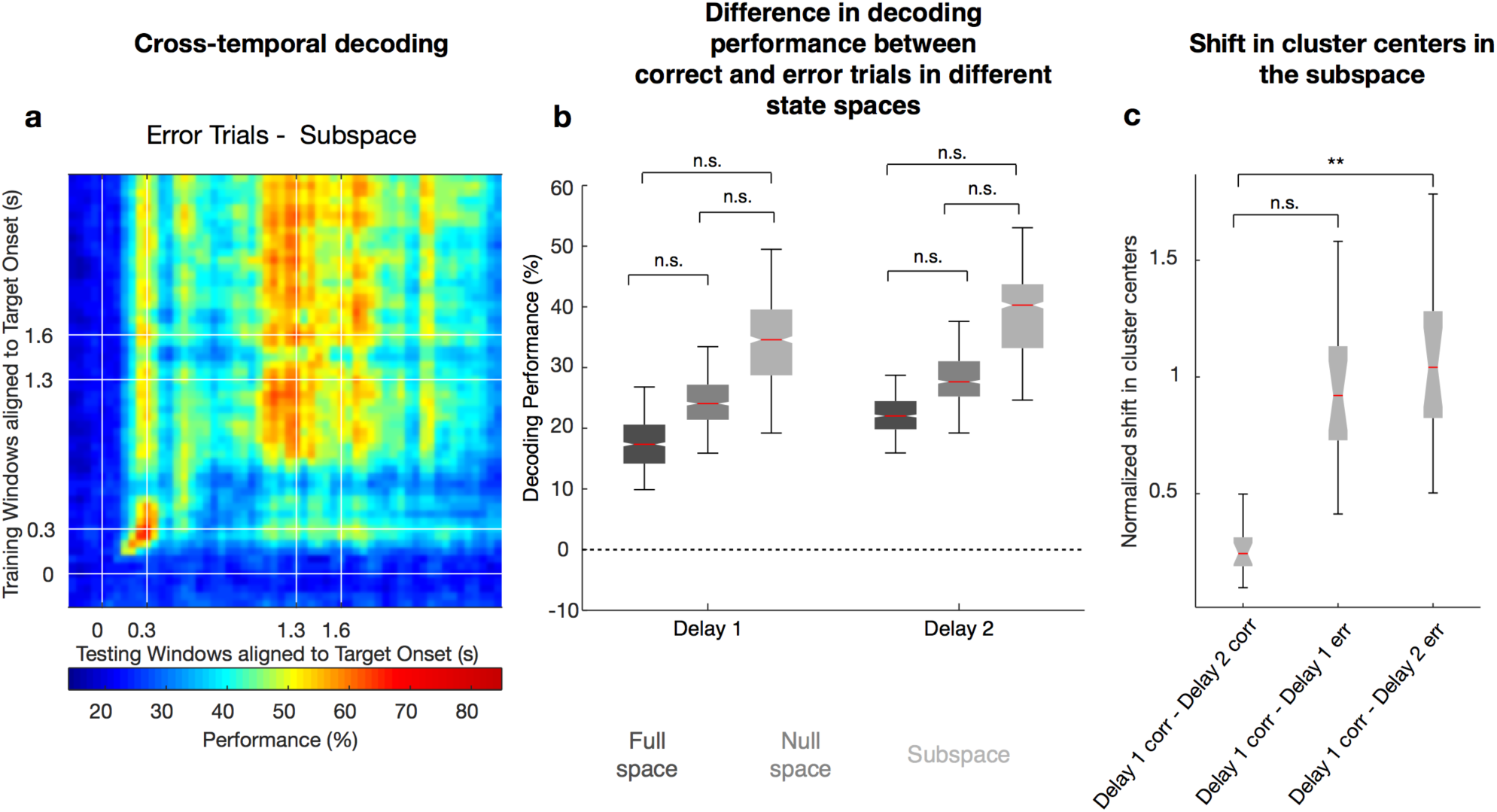
Subspace decoding in error trials. **a**, Heat map showing the cross-temporal population-decoding performance in the subspace for error trials (only 4 target locations with high error rates are included). **b**, The differences in decoding performance between correct and error trials are shown in the full space (black box-plot), subspace (gray box-plot), and the null space (dark gray box-plot) for both Delays 1 and 2. **c**, The normalized shift in cluster centers in the subspace is shown for correct trials between Delay 1 to Delay 2 (black box-plot), between correct and error trials in Delay 1 (dark gray box-plot), and between correct trials in Delay 1 and error trials in Delay 2 (gray box-plot).

While some errors may be driven by failures of working memory, as reflected in the information decay within the subspace, other factors may also lead to errors, such as failures in motor preparation or alertness. Presumably, these other factors would be reflected in information decay in spaces orthogonal to the memory subspace (i.e. the null space). In order to test this hypothesis, we measured the changes in decoding performance in both the subspace and the null space. As hypothesized, error trials also showed decreased target information in the null space (Fig. 4b). Thus, it appeared that errors may have been driven by multiple factors, including failures in memory maintenance within the subspace, and failures in non-memory factors in the null space.

A recent study showed that stable memory information could be decoded from LPFC activity despite the complex and heterogeneous temporal dynamics of single-neuron activity (Murray et al., 2017). They applied PCA to the time-averaged delay activity across stimulus conditions to find a “mnemonic subspace” in a working memory task that did not contain intervening distractors (Murray et al., 2017). When we applied the same method to our data, we found a stable subspace prior to distractor presentation. However, after distractor presentation, the code also morphed in this subspace (Supplementary Fig. 3). Another method we considered to identify a stable subspace involved using LDA. The subspace identified using LDA could be used to read out stable memory information in both delays. However, within the LDA subspace, there were significant shifts in clusters between Delay 1 and Delay 2 projections (Supplementary Fig. 4), suggesting that the apparent stability observed with decoding would break down if the task required discrimination using finer-grained memories.

Artificial neural network models (ANNs) have been shown to replicate several properties observed in the LPFC during working memory tasks (Compte et al., 2000; Wimmer et al., 2014; Murray et al., 2017; Yang et al., 2019). However, the observation of the existence of a stable subspace in the presence of code-morphing imposes new constraints on these models. Here, we tested a number of different models to attempt to replicate the following list of observations: (1) stable memory code within Delay 1 and within Delay 2, (2) code-morphing after distractor presentation, (3) existence of non-linear mixed selective neurons, and (4) existence of a stable subspace. We compared 3 popular types of ANNs to assess whether they would display these properties: a bump attractor model (Wimmer et al., 2014), a recurrent neural network (RNN) trained to report the correct target location while maintaining code stability during Delay 1, and finally a linear subspace model (Murray et al., 2017) (Table 1). Initially, we tested the models assuming that the inputs received during distractor presentation were weaker than those used during target presentation (to emulate the low behavioral relevance of the distractor in our task). Under these input parameters, none of the models were able to replicate all the properties listed above. We hypothesized that additional non-memory inputs during the distractor presentation may be required to replicate these properties. These additional inputs could be interpreted as ascending modulatory inputs, thalamic inputs, or inputs that encoded additional information, such as movement preparation or reward expectation. Using this additional input, we found that only the bump attractor model allowed us to replicate all these properties (Fig. 5). The connections in the bump attractor model consisted of strong excitation to neighboring units, and weak inhibition to units further away (see Methods). This architecture ensured a stable code during the first delay, but did not exhibit code-morphing after the distractor presentation. However, with the addition of the non-memory inputs, code-morphing occured due to the addition of new “bumps” of activity after the distractor presentation, and the locations of these new bumps were a function of the non-memory inputs (Fig. 5b). The optimization algorithm successfully identified a stable subspace from the responses in this model (Fig. 5c,d). A range of non-memory input parameters led to code-morphing in the model, suggesting that there was some degree of flexibility in the non-memory input parameters (Supplementary Fig. 5). The model also provided a couple of testable predictions: first, that the initial bump would be maintained in Delay 2; and second, that the response fields in Delay 2 would be larger than those in Delay 1. Both predictions were corroborated in the neural data (Supplementary Fig. 6a,b).

**Fig. 5.**
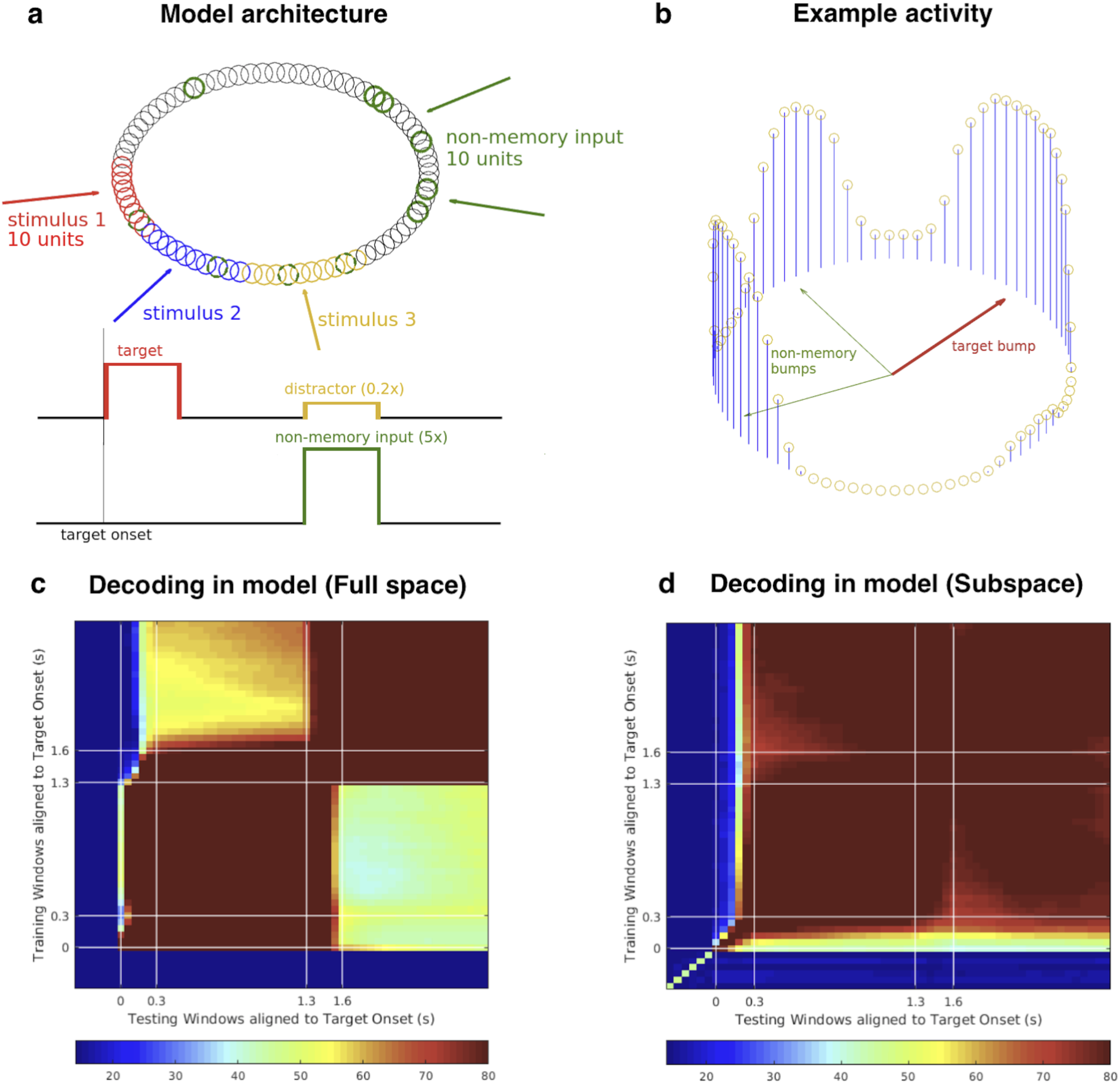
Model of code-morphing in prefrontal cortex. **a**, Bump attractor model (adapted from Wimmer et al., 2014) with 8 location inputs representing the 8 location stimuli for target or distractor, and a non-memory input that was active during the distractor presentation period. The recurrent layer contained 80 units, and each location input projected onto 10 adjacent units that were non-overlapping for different locations (red, blue, and yellow units). The non-memory input projected onto 10 random units in the population (green units), which overlapped with different location units. **b**, An example of the activity found in the recurrent layer in Delay 2 after the distractor was presented. **c**, Heat map showing the cross-temporal population-decoding performance of the model in the full space. **d**, Heat map showing the cross-temporal population-decoding result of the model after projecting into the subspace.

**Table. 1:** Comparison of different models

|  | Stable code in D1 and D2 | Code-Morphing | Existence of NMS | Existence of stable subspace |
| --- | --- | --- | --- | --- |
| Linear subspace model (Murray et al. 2016) | Yes | Yes | No | Yes |
| Trained RNN | Yes | Yes | Yes | No |
| Bump attractor model (Wimmer et al., 2014) | Yes | Yes | Yes | Yes |

Although the exact nature of the non-memory input was not clear, the model provided predictions to differentiate between different possibilities using a task with 2 consecutive distractors, separated by 1 second each (rather than the one distractor we used in the experiment). If the non-memory input corresponded to ascending modulatory or thalamic inputs that were triggered after every distractor, then code-morphing would occur after the first distractor, but not the second since the same non-memory inputs would be activated (Supplementary Fig. 6c). On the other hand, if the non-memory input corresponded to a movement preparation or reward expectation signal triggered by the stimulus that best predicted the timing of movement onset, then code-morphing would occur after the second distractor (which was closest to the movement onset), but not after the first (Supplementary Fig. 6d). These would be interesting predictions to test in future experiments.

## Discussion

Here we demonstrate that stable memory information can be read out from a population of neurons that morphs its code after a distractor is presented. This readout was enabled by a low-dimensional subspace, which was identified using an optimization algorithm that minimized the distance between projections of Delay 1 and Delay 2 activities onto state-space, while minimizing information loss. We found that an important property of neural dynamics that allowed for the existence of this subspace was the parallel movement of trajectories from Delay 1 to Delay 2 for different memory locations. Information in this subspace appears to be behaviorally relevant, since the stability breaks down in error trials. Finally, a bump attractor model replicated code-morphing and stable subspace, and revealed that code-morphing required a non-memory input during the distractor presentation period. Overall, our results show that dynamic activity in LPFC, possibly driven by non-memory inputs during distractor presentation, can be read out in a stable manner to perform a task that requires stable working memory information.

The LPFC has a high number of neurons with mixed-selective responses, which dramatically increase the dimensionality of representations (Rigotti et al., 2013). The existence of multiple subspaces within a single population of neurons may be an efficient means to use the high-dimensional activity space of brain regions (Remington et al. 2018). This view is consistent with our finding that a low-dimensional subspace within LPFC can encode stable memory information despite non-memory-related neural dynamics being present. Additional dynamics, which may reflect activity in orthogonal spaces, would allow simultaneous encoding of additional information without interfering with memory information. However, orthogonality of subspaces is not a necessity, since different types of information may interfere with each other at the cognitive and neural levels. For example, attention or movement preparation may bias or interfere with working memory information, which would suggest their encoding in subspaces which are not orthogonal to the memory subspace.

We have shown here that, in principle, a region downstream from the LPFC could read out stable memory information using the subspace. However, we have not shown that any regions are actually using this subspace to read out stable memory information. Addressing this question is particularly challenging, since even if a downstream region reads out this information, it may be immediately converted to a new task-relevant type of information, such as direction of eye movement. In order to assess whether downstream regions indeed use the memory subspace of LPFC we could assess how trial-to-trial fluctuations of population responses relate the LPFC subspace with fluctuations in downstream areas (Semedo et al., 2019). However, a more direct test would involve the manipulation of LPFC activity, either within or outside the subspace, while measuring changes in activity in downstream regions (Golub et al., 2018). To our knowledge, these experiments would not be feasible with stimulation technology available today. One approach to address this question is to generate artificial neural network models that replicate the response properties of LPFC and downstream regions, and to generate predictions of the effect of specific manipulations. Here we tested 3 different types of artificial neural network models to try to replicate the existence of a stable memory subspace in the presence of code-morphing. We found that the bump attractor model could replicate these features, while a trained randomly-connected network as well as a subspace model failed to replicate these behaviors. The model provided predictions that could be corroborated on the data, and other predictions that could not be tested with our existing data. These observations support this model as a useful abstraction of the LPFC function.

## Acknowledgements

We thank Apoorva Bhandari and Andrew Tan for discussions and suggestions on this work. This work was supported by startup grants from the Ministry of Education Tier 1 Academic Research Fund and SINAPSE to C.L., a grant from the NUS-NUHS Memory Networks Program to S.-C.Y., and a grant from the Ministry of Education Tier 2 Academic Research Fund to C.L. and S.-C.Y. (MOE2016-T2-2-117).

## Author Contributions

S.-C.Y., A.P., and C.L., conceptualized the analysis framework. A.P., C.T., and R.H. performed the analysis. L.-F.C. provided advice on the optimization technique. S.-C.Y. and C.L. guided the data analysis. All authors discussed the results, and A.P., S.-C.Y., and C.L. wrote the manuscript.

## Competing financial interests

The authors declare no competing financial interests.

### Online Methods

#### Subjects and Surgical Procedures

We used two male adult macaques (*Macaca fascicularis*), Animal A (age 4) and Animal B (age 6), in the experiments. All animal procedures were approved by, and conducted in compliance with the standards of the Agri-Food and Veterinary Authority of Singapore and the Singapore Health Services Institutional Animal Care and Use Committee (SingHealth IACUC #2012/SHS/757). The procedures also conformed to the recommendations described in Guidelines for the Care and Use of Mammals in Neuroscience and Behavioral Research (National Academies Press, 2003). Each animal was implanted first with a titanium head-post (Crist Instruments, MD, USA) before arrays of intracortical microelectrodes (MicroProbes, MD, USA) were implanted in multiple regions of the left frontal cortex (Fig. 1c). In Animal A, we implanted 6 arrays of 16 electrodes and 1 array of 32 electrodes in the LPFC, and 2 arrays of 32 electrodes in the FEF, for a total of 192 electrodes. In Animal B, we implanted 1 array of 16 electrodes and 2 arrays of 32 electrodes in the LPFC, and 2 arrays of 16 electrodes in the FEF, for a total of 112 electrodes. The arrays consisted of platinum-iridium wires with either 200 or 400 μm separation, 1 – 5.5 mm of length, 0.5 MΩ of impedance, and arranged in 4×4 or 8×4 grids. Surgical procedures followed the following steps. 24 hours prior to the surgery, the animals received a dose of Dexamethasone to control inflammation during and after the surgery. They also received antibiotics (amoxicillin 7 – 15 mg/kg and Enrofloxacin 5 mg/kg) for 8 days, starting 24 hours before the surgery. During surgery, the scalp was incised, and the muscles retracted to expose the skull. A craniotomy was performed (~ 2×2 cm). The dura mater was cut and removed from the craniotomy site. Arrays of electrodes were slowly lowered into the brain using a stereotaxic manipulator. Once all the arrays were secured in place, the arrays’ connectors were secured on top of the skull using bone cement. A head-holder was also secured using bone cement. The piece of bone removed during the craniotomy was repositioned to its original location and secured in place using metal plates. The skin was sutured on top of the craniotomy site, and stitched in place, avoiding any tension to ensure good healing of the wound. All surgeries were conducted using aseptic techniques under general anesthesia (isofluorane 1 – 1.5% for maintenance). The depth of anesthesia was assessed by monitoring the heart rate and movement of the animal, and the level of anesthesia was adjusted as necessary. Analgesics were provided during post-surgical recovery, including a Fentanyl patch (12.5 mg/2.5 kg 24 hours prior to surgery, and removed 48 hours after surgery), and Meloxicam (0.2 – 0.3 mg/kg after the removal of the Fentanyl patch). Animals were not euthanized at the end of the study.

#### Recording Techniques

Neural signals were initially acquired using a 128-channel and a 256-channel Plexon OmniPlex system (Plexon Inc., TX, USA) with a sampling rate of 40 kHz. The wide-band signals were band-pass filtered between 300 to 3000 Hz. Following that, spikes were detected using an automated Hidden Markov Model based algorithm for each channel^36^. The eye positions were obtained using an infrared-based eye-tracking device from SR Research Ltd. (Eyelink 1000 Plus). The behavioral task was designed on a standalone PC (stimulus PC) using the Psychophysics Toolbox^37^ in MATLAB (Mathworks, MA, USA). In order to align the neural and behavioral activity (trial epochs and eye data) for data analysis, we generated strobe words denoting trial epochs and performance (rewarded or failure) during the trial. These strobe words were generated on the stimulus PC, and were sent to the Plexon and Eyelink computers using the parallel port.

#### Behavioral Task

Each trial started with a mandatory period (500 ms) where the animal fixated on a white circle at the center of the screen. While continuing to fixate, the animal was presented with a target (a red square) for 300 ms at any one of eight locations in a 3×3 grid. The center square of the 3×3 grid contained the fixation spot and was not used. The presentation of the target was followed by a delay of 1000 ms, during which the animal was expected to maintain fixation on the white circle at the center. At the end of this delay, a distractor (a green square) was presented for 300 ms at any one of the seven locations (other than where the target was presented). This was again followed by a delay of 1000 ms. The animal was then given a cue (the disappearance of the fixation spot) at the end of the second delay to make a saccade towards the target location that was presented earlier in the trial. Saccades to the target location within a latency of 150 ms and continued fixation at the saccade location for 200 ms was considered a correct trial. An illustration of the task is shown in Figure 1a. One of the animals was presented with only 7 of the 8 target locations because of a behavior bias in the animal.

#### Firing rate normalization

The firing rate of each neuron (averaged across trials with 100 ms windows with 50 ms of overlap) was converted to a z-score by normalizing to the mean and standard deviation of the instantaneous firing rates from 300 ms before target onset to target onset. These z-scores were then used for the state-space and subspace analyses. Our database initially consisted of 256 LPFC neurons, but we excluded 12 neurons as they exhibited responses that were very similar to other neurons, suggesting that they were the result of over-clustering during spike sorting.

#### Subspace Identification

We used the following optimization equation to identify the subspace:

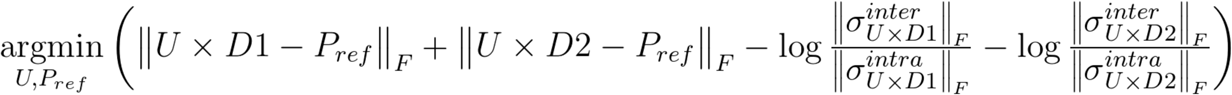

We postulated that there was a matrix transformation, *U*, that would be able to transform both the Delay 1 (*D1*) and Delay 2 (*D2*) responses to a subspace where the distance between the corresponding target responses in the two delays were minimized. In order to avoid the trivial solution where *U* was all zeros, which meant that all points were transformed to the origin, and thus have minimal distance between *D1* and *D2* responses, we performed the optimization by computing the distance between *U x D1* and a set of points *P*_*ref*_, between *U x D2* and the same set of points, *P*_*ref*_, and minimizing the sum of those two distances. *P*_*ref*_ can be thought of as the ideal end points of the transformation, so this meant that we should be able to minimize the distance if *U x D1, U x D2*, and *P*_*ref*_ ended up being the same points. In practice, we first reduced the normalized firing rates from 800 ms to 2500 ms after target onset (i.e. 500 ms before the end of *D1* to the end of *D2*) of the 244 LPFC neurons to a smaller number of Principal Components Analysis (PCA) components that accounted for 90% of the variance, which turned out to be 58. We subsequently used these 58 dimensions to be the full space in the remainder of this paper. We then took 50 responses for each of the 7 target locations to create *D1* and *D2* matrices that were 58×350 in size, where each of the columns was the averaged z-score for the 244 cells over the last 500 ms of *D1* or *D2* for one target location projected onto the 58 components. *U* was then a *u*_*dim*_x58 matrix, while *P*_*ref*_ was a *u*_*dim*_x350 matrix, and both were initialized with random values.

We also wanted the post-transformation distances between the responses to different target locations to be maximized to retain as much information as possible. As such, we added the two terms at the end to maximize the ratio of the average inter-cluster (*σ*^*inter*^) to the intra-cluster (*σ*^*intra*^) distances after the transformation.

The optimization was performed using the “fmincon” function in Matlab to minimize the cost function shown above using the sequential quadratic programming (sqp) algorithm. There were no additional constraints imposed on this optimization. The results shown in Figures 2–4 contain results from an optimized 8-dimensional subspace (i.e. *u*_*dim*_ = 8). We also performed this optimization to yield up to a 10-dimensional subspace, and found that the subspace was not qualitatively different for 7, 8, 9, or 10 dimensions.

Using the *U* matrix returned by the optimization, we concatenated *U x D1* and *U x D2* together to create a matrix that was 8×700, and then performed PCA again to obtain the top two components (SP1 & SP2) used to plot the points in the subspace in Figure 2b, as well as the other components used in Figure 2f.

In order to compute the contribution of each neurons, we took the 8×58 *U* matrix, and multiplied it by the 58×244 PCA components to obtain a 8×244 weight matrix. We computed the magnitude for all the weights in the matrix, and then normalized them by the largest weight. We then computed the average weight for each of the neurons by taking the average for each column. Neurons were identified as NMS, LMS, and CS based on a 2-way ANOVA with independent variables for target location and trial epoch, as described in Parthasarathy et al., 2017).

#### Cross-Temporal Decoding

In order to assess the stability of the population code, we used data at each time point to train a decoder based on Linear Discriminant Analysis (LDA), built using the classify function in MATLAB, and tested the decoder on data from other time points as described in Parthasarathy et al. (2017). One minor difference in the current work is that instead of identifying separate PCA components for D1 and D2, we used the same 58 PCA components that were used for the subspace analysis described above, which spanned both D1 and D2.

In order to compute the decoding performances in Figure 2d, we averaged the cross-temporal performance for classifiers trained in the last 500 ms of Delay 1 (800–1,300 ms after target onset) and tested on data from Delay 1 (labelled as *LP*_*11*_) and Delay 2 (1,800–2,300 ms after target onset, labelled as *LP*_*12*_). Similarly, we averaged the cross-temporal performance for classifiers trained in Delay 2 and tested on data from Delay 2 (*LP*_*22*_) and Delay 1 (*LP*_*21*_). This then allowed us to quantify the change in performance when decoding across delay periods. By using different subsets of the data for training and testing, we obtained distributions of performance accuracies that we were able to use for testing statistical significance (described below).

#### State Space Analysis

After projecting the responses in D1 and D2 into the 58-component PCA space, we computed the center of each of the 7 clusters representing target locations in D1 and D2. We then computed the inter-delay distance between the centers of corresponding target locations, and used that to generate the values for the full space plotted in Figure 2e. We also computed the distance between corresponding target locations in D1 and during the presentation of the distractor. As a control, we divided both delay periods into an early 250 ms and a late 250 ms, and computed an equivalent intra-delay distance. In order to account for the variability within each cluster, we computed the average intra-cluster distances for all 14 clusters over 1000 bootstrapped samples. The intra-cluster distances were used to normalize both the inter-delay and intra-delay distances. By using different subsets of the data to form D1 and D2, we obtained distributions of both inter-delay and intra-delay distances that we were able to use for testing statistical significance (described below).

After transforming D1 and D2 into the subspace using the *U* matrix, we repeated the steps described above to generate the values for the subspace plotted in Figure 2e.

#### Trajectory Directions

After projecting the responses in D1 and D2 into the 58-component PCA space, we computed the 7 vectors that connected target locations in D1 with the corresponding locations in D2. We then used a measure known as phase locking value (PLV, Lachaux et al., 1999) to quantify the similarity between the vectors. Briefly, this measure averaged the 7 vectors together and computed the magnitude of the average vector. If the vectors were very similar, the PLV will be close to 1. If the vectors were quite dissimilar, the magnitude of the average vector will be close to 0. For comparison, we shuffled the target locations in D2, and re-computed the 7 vectors before computing the PLV. By using different subsets of data to form D1 and D2, we obtained distributions of PLVs that we were able to use for testing statistical significance (described below).

#### Null Space

The *U* matrix described above transformed the *D1* and *D2* vectors in the full space to the *SP1* and *SP2* vectors in the subspace. We then identified a null space where a basis set of vectors, *v*, returned *U(v) = 0*. This meant that the null space consisted of responses in the full space that were not captured in the subspace. For this, we used the “null” function in Matlab to compute the null space given the 8×58 *U* matrix. This returned a 50×58 *V* matrix that defined the null space.

#### Statistics

We considered two bootstrapped distributions to be significantly different if the 95th percentile range of the two distributions did not overlap. We also computed an estimated p-value for this comparison using the following formula (Ojala et al., 2010),

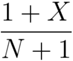

where X represents the number of overlapping data points between the two distributions and N represents the number of bootstraps. With this computation, and the N = 1000 bootstraps we used throughout the paper, two distributions with no overlap will result in a p-value < 0.001, and two distributions with x% of overlap will result in a p-value ~ x/100.

In addition to the estimated p-value, we also computed the effect size of the comparison using a measure known as Hedges’ g, computed using the following formula (Hedges 1981),

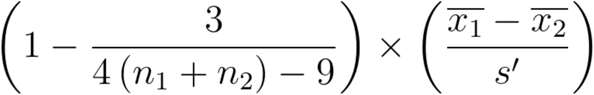

where

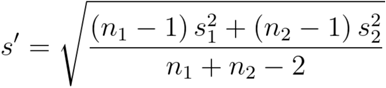

refers to the mean of each distribution, n refers to the length of each distribution, and s refers to the standard deviation of each distribution.

No statistical methods were used to pre-determine sample sizes, but our sample sizes are similar to those reported in previous publications. The majority of our analyzes made use of non-parametric permutation tests, and as such, did not make assumptions regarding the distribution of the data. No randomization was used during the data collection, except in the selection of the target and distractor locations for each trial. Randomization was used extensively in the data analyzed to test for statistical significance. Data collection and analysis were not performed blind to the conditions of the experiments. No animals or data points were excluded from any of the analyses. Please see additional information in the Life Sciences Reporting Summary.

#### Model

In order to replicate the features we found in our neural data, we tested three types of artificial neural network models that are typically used to model working memory with persistent activity: 1) linear subspace model, 2) trained RNN model with backpropagation, and 3) bump attractor model. We found that only the bump attractor model was able to replicate all the significant features found in our data. For the bump attractor model, we used *N* = 80 firing-rate units as the whole population, and used simplified discrete time equations to describe the dynamics of the population:

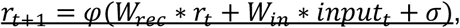

where *r*_*t*_ was the population firing rate at time *t*, 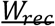 was the recurrent connection weight between units, 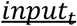was the external input at time *t*, 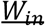 was the loading weight of input signal to the population, *σ* ~ *N*(0,0.1) was a noise term, and *φ*(*x*) was a piecewise nonlinear activation function adopted from [Compte 2014]:

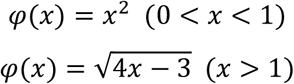

The matrix, 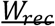, had a diagonal shape with stronger positive values near the diagonal, and weaker negative values elsewhere, such that only a few neighboring units were connected via excitatory weights to each other while being connected via inhibitory weights to the rest. In this way, a structured input signal to adjacent units was able to generate a local self-sustaining bump of activity. There were eight input units, representing the eight spatial target locations in the animal’s task. For each input unit, the loading weight matrix, *W*_*in*_, specified 10 adjacent units in the population to receive the input signal, and the loading population for each input unit were non-overlapping (Fig. 5a, different colors of stimuli).

We tested a range of distractor activity levels relative to the target, and found that higher distractor activity led to higher distractor decoding accuracy, as expected. We also found that in this model, with distractor activity levels comparable to the target, target decoding performance exhibited only a small decrease. However, distractor activity on its own did not generate code-morphing. The red circles indicates a range of distractor activity levels that replicated lower distractor decoding performance compared to target decoding performance (with a ratio of ⅓) as reported in Parthasarathy et al. (2017). Within this range, we chose 0.2 as the distractor activity level. We also randomly chose *n* individual units from the whole population as ‘non-memory’ units (Fig. 5a, green circles). In each simulated trial, the *n* ‘non-memory’ units received inputs during the distractor period with strength, *s*. We tested different pairs of combinations of *n* and *s*, and found that the pairs that successfully replicated code-morphing exhibited an anti-correlation with a range of flexibility, as depicted in the red circles in Supplementary Figure 5a,b.

## Data Availability

The data that support the findings of this study are available from the corresponding authors upon reasonable request.

## Supplementary Figures

**Supplementary Figure 1.**
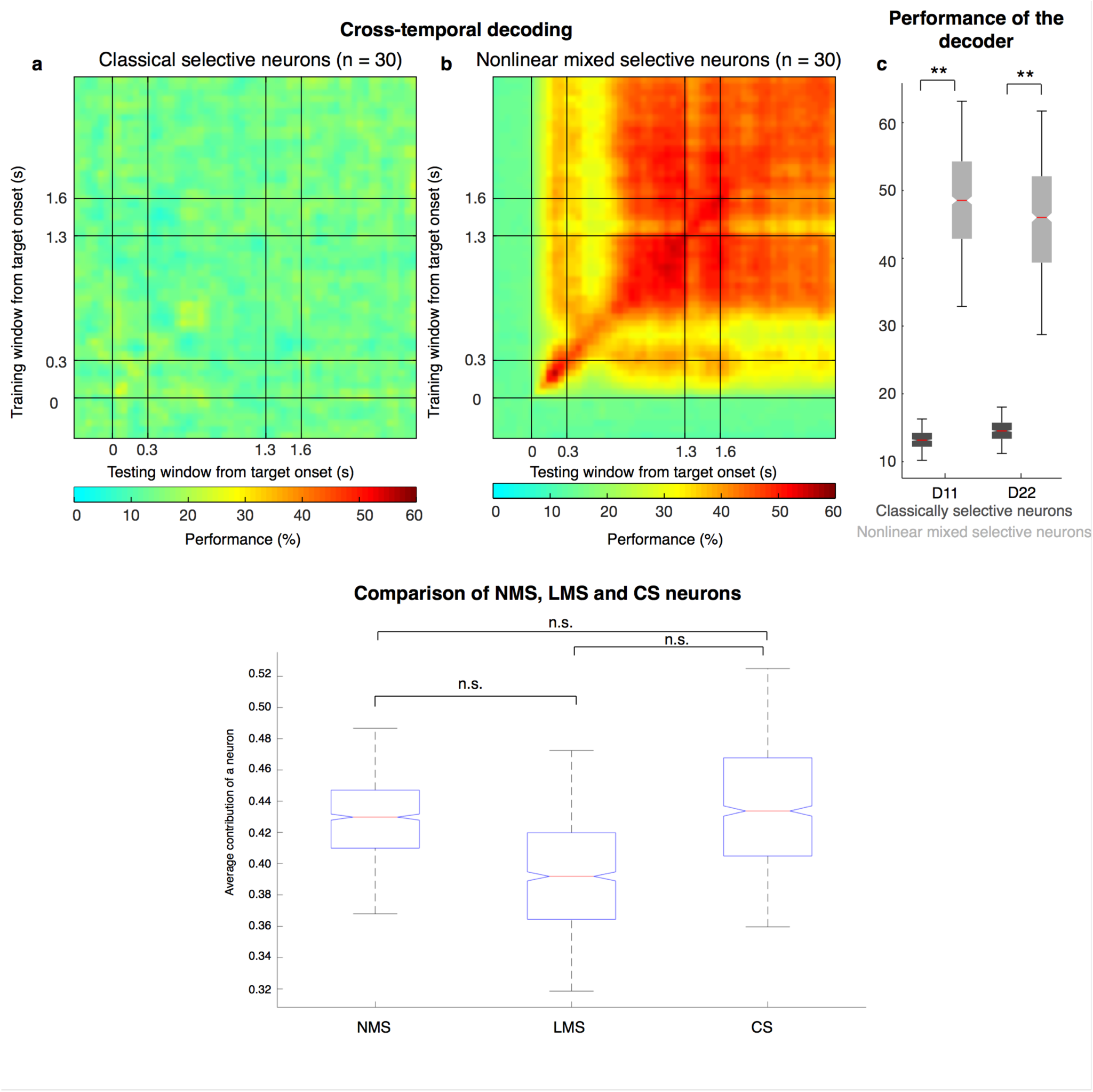
Subspace decoding using subpopulations of LPFC neurons. **a**, Heatmap showing subspace decoding using only CS neurons. **b**, Heatmap showing subspace decoding using only NMS neurons. **c**, Box-plots showing the performance of the subspace decoder for CS (black box-plots) and NMS neurons (gray box-plots) in Delays 1 and 2. The performance for the CS neurons were significantly lower than those for the NMS neurons (D11: P < 0.001 g = 15.27 D22: P < 0.001 g = 14.39). **d**, Contribution of different types of neurons (including neurons with linear mixed-selectivity, LMS) to the subspace shown in Figure 2b. There were no significant differences between the 3 distributions (P ≈ 0.76 for NMS and, LMS, P ≈ 0.73 for LMS and, CS and P ≈ 0.89 for NMS and CS).

**Supplementary Figure 2.**
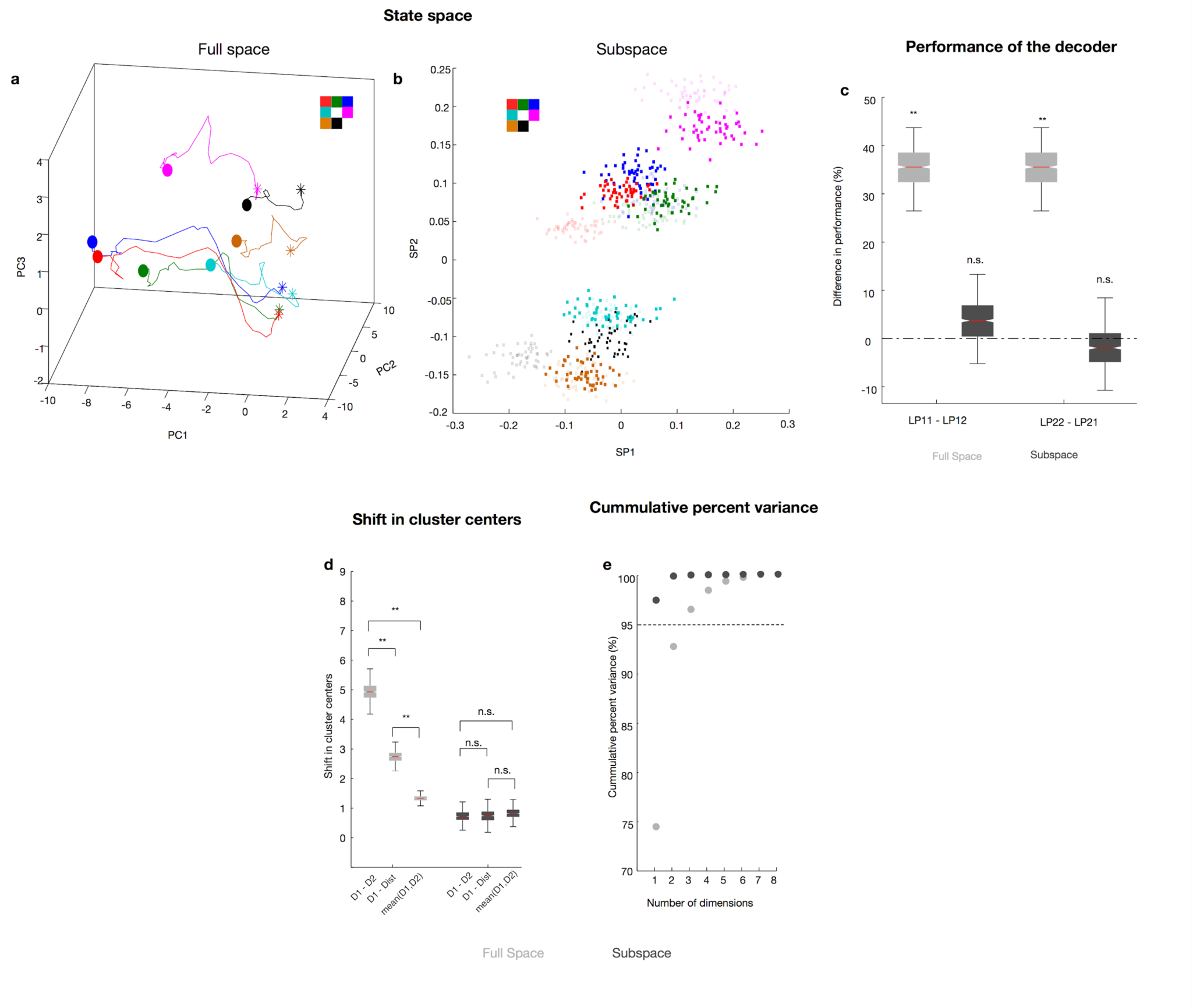
Characterization of the full space and subspace found using only NMS neurons. **a**, Responses of the NMS neurons when projected onto the first three principal components (PC) of the combined Delay 1 and Delay 2 response space. The responses for different target locations are color-coded using the color scheme shown in the top left. The trajectories illustrate the evolution of the responses from 500 ms before the end of Delay 1 (square), distractor onset (first triangle), distractor offset (second triangle), and the end of Delay 2 (circle). The trajectories of only 4 of the 7 target locations are shown here for clarity. **b**, Delay 1 (plotted using crosses) and Delay 2 (plotted using squares) responses after projection into the top 2 PCs of the subspace. Points for different target locations are color-coded according to the color scheme shown in the top left. **c**, The difference in decoding performance when a decoder that was trained on Delay 1 responses was tested on Delay 1 (LP11) or Delay 2 (LP12) responses are shown in the gray box-plot labelled LP11 - LP12, and vice versa (gray box-plot labelled LP22 - LP21, P < 0.001 for LP11 - LP12 and LP22 - LP21). The equivalent performance differences in the subspace are shown in the black box-plots (P ≈ 0.45 for LP11 - LP12 and P ≈ 0.66 for LP22 - LP21). **d**, The shift in cluster centers from Delay 1 to Delay 2 (labeled D1 - D2) averaged across target locations, from Delay 1 to the distractor presentation period (D1 - Dist), and the intra-delay shifts in both Delays 1 and 2 (intra-delay) are shown in the full space (gray box-plot, P < 0.001 for D1 - D2 and D1 - Dist, g = 20.75 for D1 - D2 and 16.54 for D1 - Dist), and in the subspace (black box-plot, P ≈ 0.97 for D1 - D2 and P ≈ 0.96 for D1 - D2). **e**, The cumulative explained variance is plotted as a function of the number of PCs for the full space (plotted in gray) and the subspace (plotted in black).

**Supplementary Figure 3.**
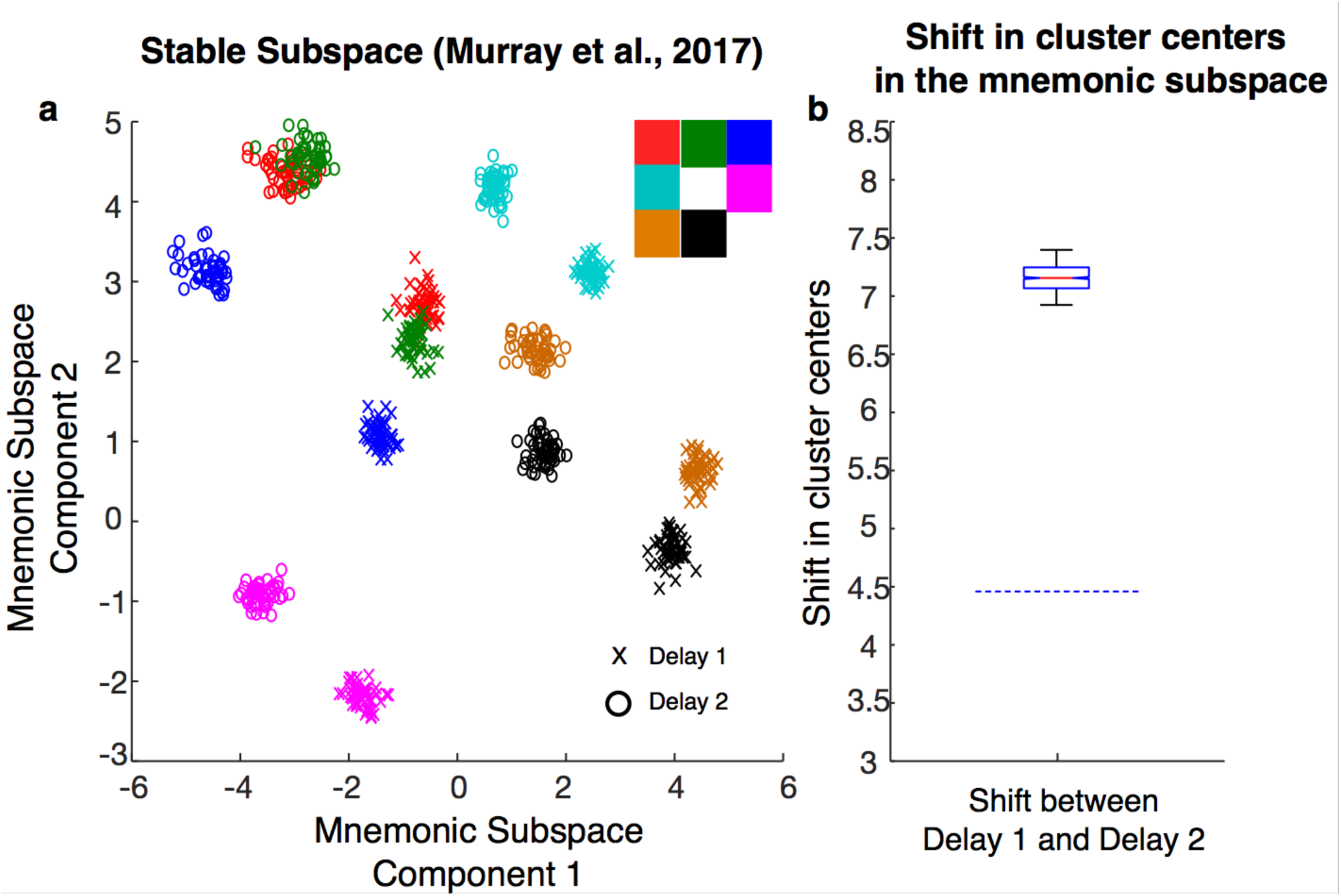
Mnemonic Subspace. **a**, Responses of LPFC neurons when projected onto the first two principal components of the mnemonic subspace. The responses for different target locations are color-coded using the color scheme shown in the top right. Responses in Delay 1 are plotted using crosses (x), while responses in Delay 2 are plotted using circles (o). **b**, The shift in cluster centers from Delay 1 to Delay 2 (averaged across target locations) are shown for the mnemonic subspace (computed over 58 components, which accounted for 95% of the variance). The dotted lines show the 97.5th percentile of the intra-delay shifts averaged across target locations. The inter-delay shifts were significantly different from the intra-delay shifts (P < 0.001, g = 31.62).

**Supplementary Figure 4.**
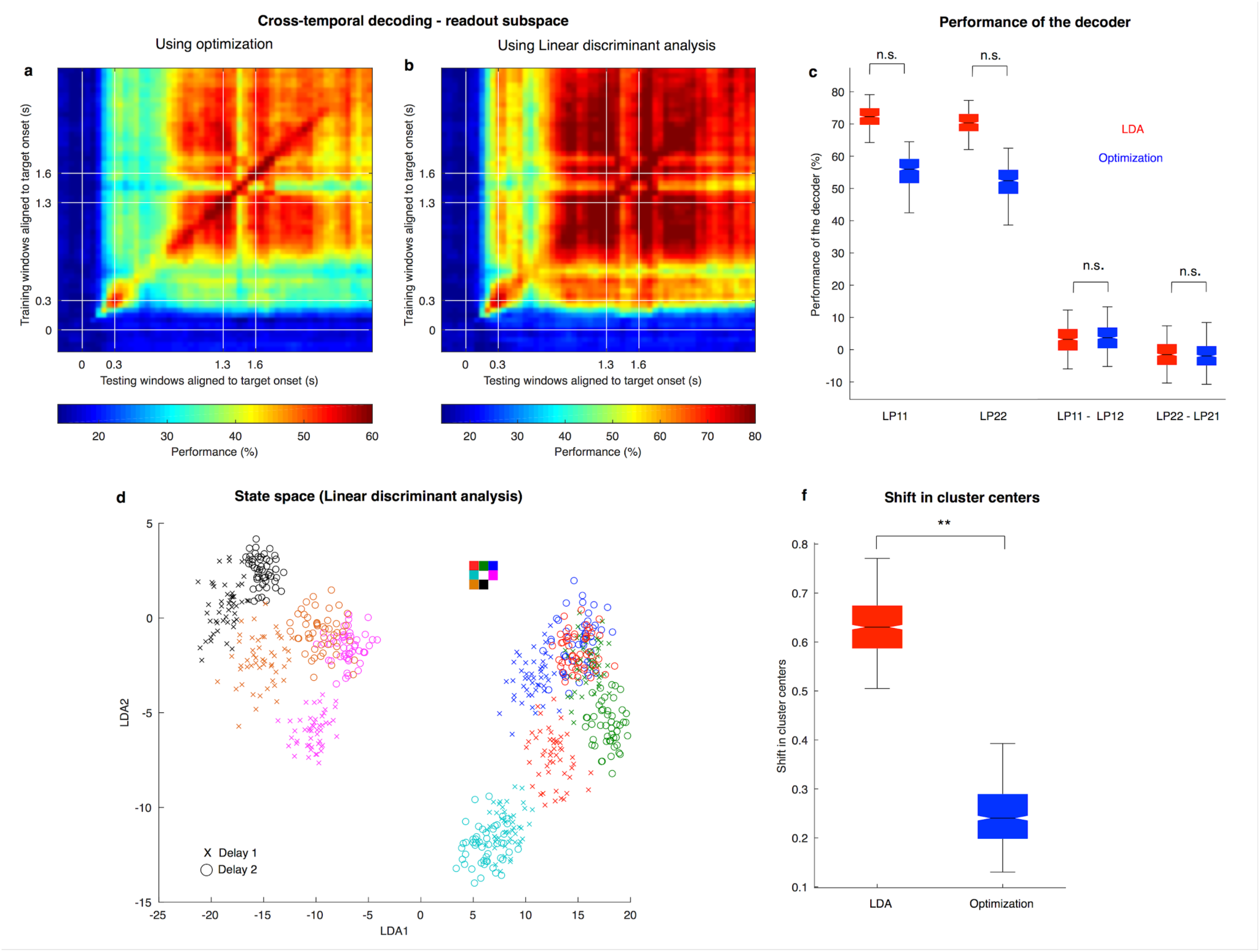
LDA Subspace. **a**, Heat map showing the cross-temporal population-decoding performance after the population responses were projected onto the optimized subspace (shown previously in Fig. 2c). **b**, Heat map showing the cross-temporal population-decoding performance after the population responses were projected onto the LDA subspace. **c**, The decoding performance in the LDA subspace was not significantly different from the optimized subspace (LP11: P ≈ 0.13; LP22: P ≈ 0.14; LP11 - LP12: P ≈ 0.99; LP22 - LP21: P ≈ 0.98). **d**, Delay 1 (plotted using crosses, x) and Delay 2 (plotted using circles, o) responses after projecting into 2 of the LDA boundaries. Points for different target locations are color-coded according to the color scheme shown in the top right. **e**, The cumulative explained variance is plotted as a function of the number of PCs for the LDA subspace (plotted in red) and the optimized subspace (plotted in blue). **f**, The shift in cluster centers from Delay 1 to Delay 2 (averaged across target locations) are shown for the LDA subspace (plotted in red, computed over 8 components, which accounted for 95% of the variance) and the optimized subspace (plotted in blue, shown previously in Fig. 2e). The shift in the LDA subspace was significantly larger than that in the optimized subspace (P ≈ 0.01, g = 9.07).

**Supplementary Figure 5.**
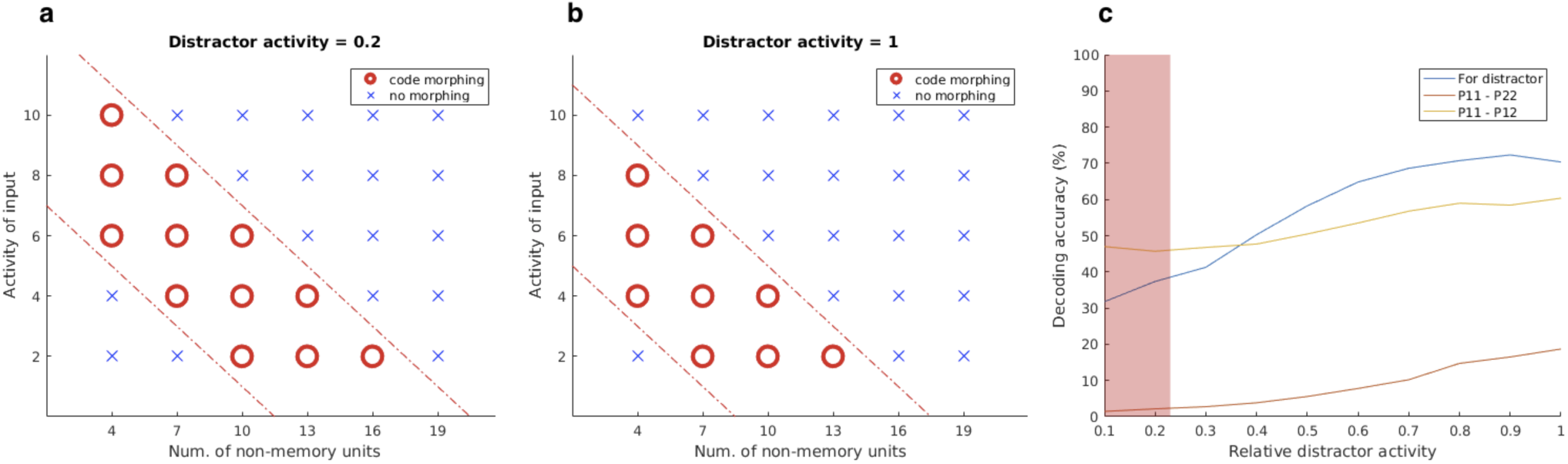
Parameters in the bump attractor model. **a**,**b**, We tested different pairs of combinations of different numbers of non-memory units, *n*, and activity level of the non-memory inputs, *s*, along with the activity level of the distractor (0.2 in **a**, and 1.0 in **b**), and found that the pairs that successfully replicated code-morphing exhibited an anti-correlation between *n* and *s*. **c**, The decoding performance in the model for the distractor location (plotted in blue) is shown for different distractor activity levels relative to the target activity level. The difference in performance between *LP*_*11*_ and *LP*_*22*_, and between *LP*_*11*_ and *LP*_*12*_, are plotted in red and yellow, respectively.

**Supplementary Figure 6.**
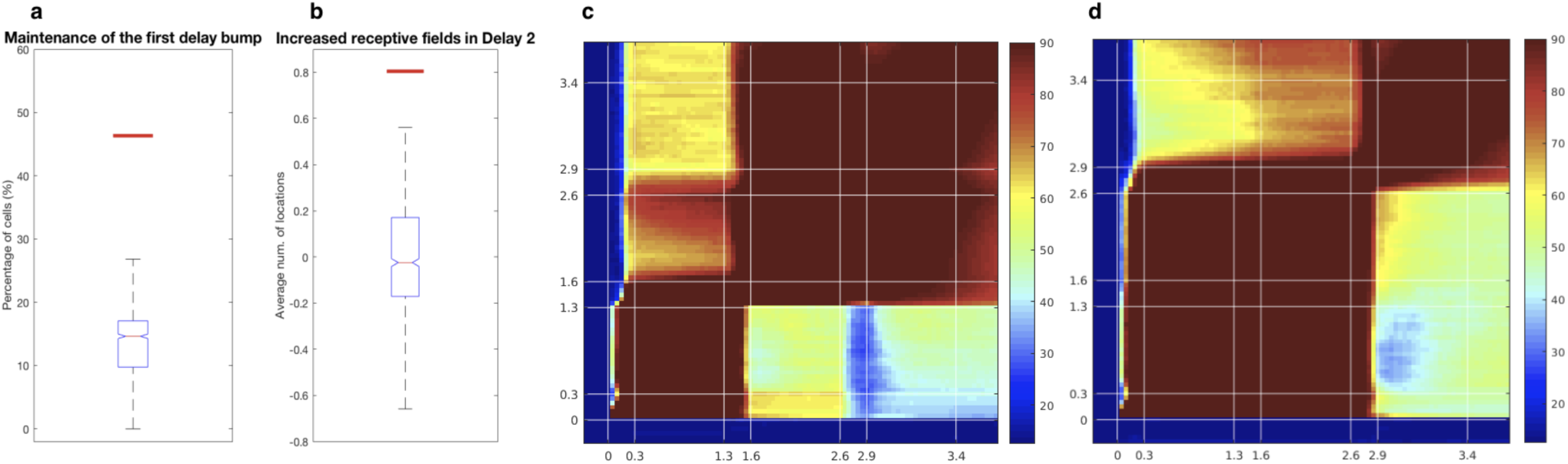
Predictions of the bump attractor model. **a**, The percentage of NMS cells that maintained its selectivity in Delay 1 into Delay 2 (characterized simply by a lack of change in the most responsive location in both delays) is shown in the red horizontal line, while the box-plot illustrate the 5th and 95th percentile of the null distribution, which shuffled the responses for different locations within each delay (*P* < 0.001, *g* = 5.85) **b**, The number of responsive locations for each neuron increased in Delay 2 compared to Delay 1 (*P* < 0.001, *g* = 3.26). This corresponded to the addition of the non-memory bump to the target location bump in Delay 2. The red horizontal line shows the average increase in locations, while the box-plots illustrate the 5th and 95th percentile of the null distribution, which shuffled responses for different locations between the two delays. **c**, In a simulation with 2 distractors presented, the non-memory input (assumed to be an ascending input) was activated during both distractor presentations, so code morphing only occurred after the first distractor was presented. **d**, In a simulation with 2 distractors presented, the non-memory input (assumed to be something like movement preparation or reward expectation) was only activated during the second distractor presentation, so code morphing only occurred after the second distractor was presented.

**Supplemental Movie 1.** LPFC and FEF state space movie

http://www.dropbox.com/s/9q5n5577uo5602m/StateSpaceCombined.mp4?dl=0

